# Beyond power: A large-scale characterization of intrinsic brain oscillatory activity

**DOI:** 10.64898/2026.05.12.724595

**Authors:** Enrique Stern, Almudena Capilla

**Affiliations:** Departamento de Psicología Biológica y de la Salud, Facultad de Psicología., Universidad Autónoma de Madrid, Madrid, Spain

**Keywords:** Brain oscillations, Human, Magnetoencephalography (MEG), Power spectrum, Resting state

## Abstract

Most of what we currently know about brain oscillations is derived from Fourier-based spectral power methods. While widely adopted, these procedures introduce methodological limitations and inherently overlook informative neurophysiological features that often remain unreported. In this study, we characterized resting-state oscillatory activity from human magnetoencephalography (MEG) recordings (N = 128) by integrating two complementary approaches. First, oscillatory episodes were detected at the source level with sBOSC. Subsequently, the ByCycle algorithm was applied to these episodes to extract individual cycle features. Results revealed that the brain engages in oscillatory activity for only ∼25% of the recording time, with occipital and parietal regions accounting for the highest temporal prevalence across canonical frequency bands. Furthermore, oscillatory episodes lasted an average of 4.6 cycles, reinforcing the view of neural oscillations as transient bursts. Region-specific duration and power measures revealed distinct anatomical organizations offering complementary physiological information. Finally, by extracting the instantaneous amplitude, period, and waveform asymmetry of individual cycles, we successfully dissociated sinusoidal occipital alpha waves from the asymmetric sensorimotor mu rhythm. By moving beyond traditional power-centric analyses, this approach provides a comprehensive characterization of spontaneous oscillatory activity, thereby offering new insights into the spatial, temporal, and spectral structure of human brain oscillations.

## 1. Introduction

Despite nearly a century of electrophysiological research (Berger, 1929), the fundamental principles underlying brain oscillations have yet to be fully understood. Over the years, these rhythmic patterns of electrophysiological activity have been associated with multiple cognitive functions (Buzsáki, 2002; Cannon et al., 2014; Herweg et al., 2020; Klimesch, 2012; Lopes da Silva, 2013; Staudigl & Hanslmayr, 2013; Thut et al., 2012), brain states (Baker et al., 2014; Chen et al., 2020; Seedat et al., 2020; Van Es et al., 2025), and a wide range of clinical conditions (Buzsáki & Watson, 2012; Schnitzler & Gross, 2005). Contemporary neuroscience has increasingly shifted away from relegating neural oscillations to mere epiphenomena, focusing instead on establishing their casual role in cognition (Herrmann et al., 2016; Singer, 2018; Singer & Effenberger, 2025). In this context, multiple theoretical frameworks have emerged, positing neural oscillations as a fundamental mechanism of brain communication (Bonnefond et al., 2017; Buzsáki, 2006; Fries, 2005, 2015; Palva & Palva, 2018; Vinck et al., 2023; Wang, 2010).

In electrophysiological studies of oscillatory activity, three main parameters are commonly used to characterize the signal (Donoghue et al., 2022; Keil et al., 2022; Vinck et al., 2023): frequency, defined as the number of cycles per second (Hz); power, defined as the magnitude of activity within a specific frequency; and phase, defined as the position of the oscillatory cycle at a given moment. However, our current understanding of neural oscillations is mostly based on power measures derived from Fourier-based methods. While widely adopted, several caveats arise from this procedure (Donoghue et al., 2020; Gross, 2014; Gyurkovics et al., 2021, 2022).

First, electrophysiological brain signals comprise both oscillatory and aperiodic (1/f-like) activity. However, standard Fourier-derived power spectra mathematically conflate these components. Consequently, an increase in spectral power does not unequivocally indicate the presence of an oscillation (Donoghue et al., 2020; Gerster et al., 2022). To address this limitation, several methods have recently emerged to isolate genuine oscillatory activity (Brady & Bardouille, 2022; Donoghue et al., 2020; Kosciessa et al., 2020; Seymour et al., 2022; Stern et al., 2026; Wen & Liu, 2016).

Beyond the conflation of signal components, conventional Fourier-based approaches are inherently constrained by temporal windowing. With these methods, phase and power estimates are smoothed approximations calculated over fixed or adaptive temporal windows that scale with frequency (Cohen, 2019; Daubechies, 1990). In contrast to instantaneous phase and power measures (Cohen, 2014; Huang et al., 1998), this approach introduces temporal smearing, thereby compromising temporal resolution, and potentially underestimating the true magnitude of an oscillation.

This loss of temporal precision is frequently exacerbated by standard analytical practices. Although not mandatory, a conventional practice in cognitive neuroscience experiments is to average power spectra across multiple trials to enhance the signal-to-noise ratio (SNR) (Gross, 2014). This often leads to the misinterpretation of short-lived, transient events as sustained activity, as an effect of averaging non-negative power values (Jones, 2016; Tal et al., 2020; Van Ede et al., 2018).

Finally, while Fourier-based spectral decomposition procedures assume purely sinusoidal waveforms, brain oscillations often exhibit non-sinusoidal morphologies (Cole et al., 2017; Cole & Voytek, 2017). Forcing a non-sinusoidal signal into a sinusoidal representation inevitably results in the generation of spurious harmonics, which may be misinterpreted as genuine, independent oscillations (Lozano-Soldevilla et al., 2016; Schaworonkow, 2023). Furthermore, because Fourier-based methods decompose signals using sinusoidal basis functions, they fail to capture waveform morphology, a feature hypothesized to carry relevant neurophysiological information (Cole & Voytek, 2017; Jones, 2016).

To address these methodological challenges, we characterized intrinsic human brain oscillatory activity using magnetoencephalography (MEG) data from 128 participants of the Open MEG Archive (OMEGA) database (Niso et al., 2016) by integrating two complementary methods. Because oscillations must first be disentangled from non-oscillatory activity, our analysis was built upon oscillatory episodes previously identified using sBOSC (i.e., source-Better OSCillation detection) (Stern et al., 2026), a variant of the BOSC family (Kosciessa et al., 2020; Seymour et al., 2022; Whitten et al., 2011) specifically designed to detect genuine oscillatory events at the source level. Focusing on those segments where oscillations were present, we then applied ByCycle, an algorithm that quantifies the amplitude, period (i.e., the inverse of frequency), and asymmetry of oscillations on a cycle-by-cycle basis (Cole & Voytek, 2019). By identifying oscillatory events and adopting the oscillatory cycle as the fundamental unit of analysis, we can address critical neurophysiological questions that remain obscured by conventional Fourier-based approaches. What proportion of time does the brain engage in genuine oscillatory activity? Are these oscillations sustained over prolonged periods, or do they primarily manifest as short-lived, transient bursts? In which brain regions do oscillations deviate from standard sinusoidal waveforms?

In the present study, we aimed to answer these questions by systematically characterizing the distribution, duration, and waveform morphology of intrinsic neural oscillations. We hypothesized that brain regions are not in a perpetual state of rhythmicity, but instead manifest intermittent bouts of oscillatory activity. Furthermore, we anticipated a spatial dissociation of oscillatory activity, whereby distinct brain regions exhibit characteristic spectral profiles and waveform features.

## 2. Methods

To extract individual cycle metrics, we implemented a sequential two-step process that applies two algorithms. First, to detect oscillatory cycles in the MEG signal, we used sBOSC, a method previously developed by our group to identify genuine oscillations at the source-level (Stern et al., 2026). Subsequently, to quantify individual cycle measures we employed ByCycle, a complementary analysis that extracts specific waveform features operating within the time domain (Cole & Voytek, 2019). We briefly introduce the fundamental principles of sBOSC and ByCycle in the following sections.

The full code required to reproduce the method, analyses, and figures in this work is openly available at https://github.com/necog-UAM/.

### 2.1. Data acquisition

Anonymized resting-state MEG recordings (275 axial gradiometers and 26 reference sensors, 2400 Hz sampling rate, 600 Hz low-pass anti-aliasing filter) and T1-weighted Magnetic Resonances Images (MRIs) were retrieved from the OMEGA database (Niso et al., 2016). Data were acquired at the Montreal Neurological Institute (MNI, McGill University), in accordance with the Declaration of Helsinki and with approval from the institutional ethics committees. MEG recordings were acquired in an upright position during a relaxed, eyes-open wakeful state. In this study, we analyzed data from 128 healthy participants (68 males, 118 right-handed, mean age 30.5 ± 12.4 [M ± SD], range 19-73 years), previously preprocessed in an earlier work. In brief, MEG signals were denoised, high-pass filtered at 0.05 Hz, and cleaned using independent component analysis (ICA), with additional removal of line noise and artifact-contaminated segments (see Capilla et al., 2022 for further details).

### 2.2. sBOSC

sBOSC is an oscillatory detection algorithm designed to identify neural oscillatory episodes directly at their brain sources (Stern et al., 2026). Briefly, it consists of the following steps.

First, source-level activity was reconstructed using linearly constrained minimum variance beamforming (LCMV; Van Veen et al., 1997) (Figure 1a). Co-registration of each participant’s MRI with the MEG head coordinate space was obtained from Capilla et al. (2022) (https://github.com/necog-UAM/OMEGA-NaturalFrequencies/tree/main/coregistration). Lead fields were computed on a 1-cm grid using a realistic single-shell volume conduction model (Nolte, 2003). Spatial filter weights were then estimated from the covariance of artifact-free data with 10% regularization and applied to reconstruct source-level time series.

**Figure 1.**
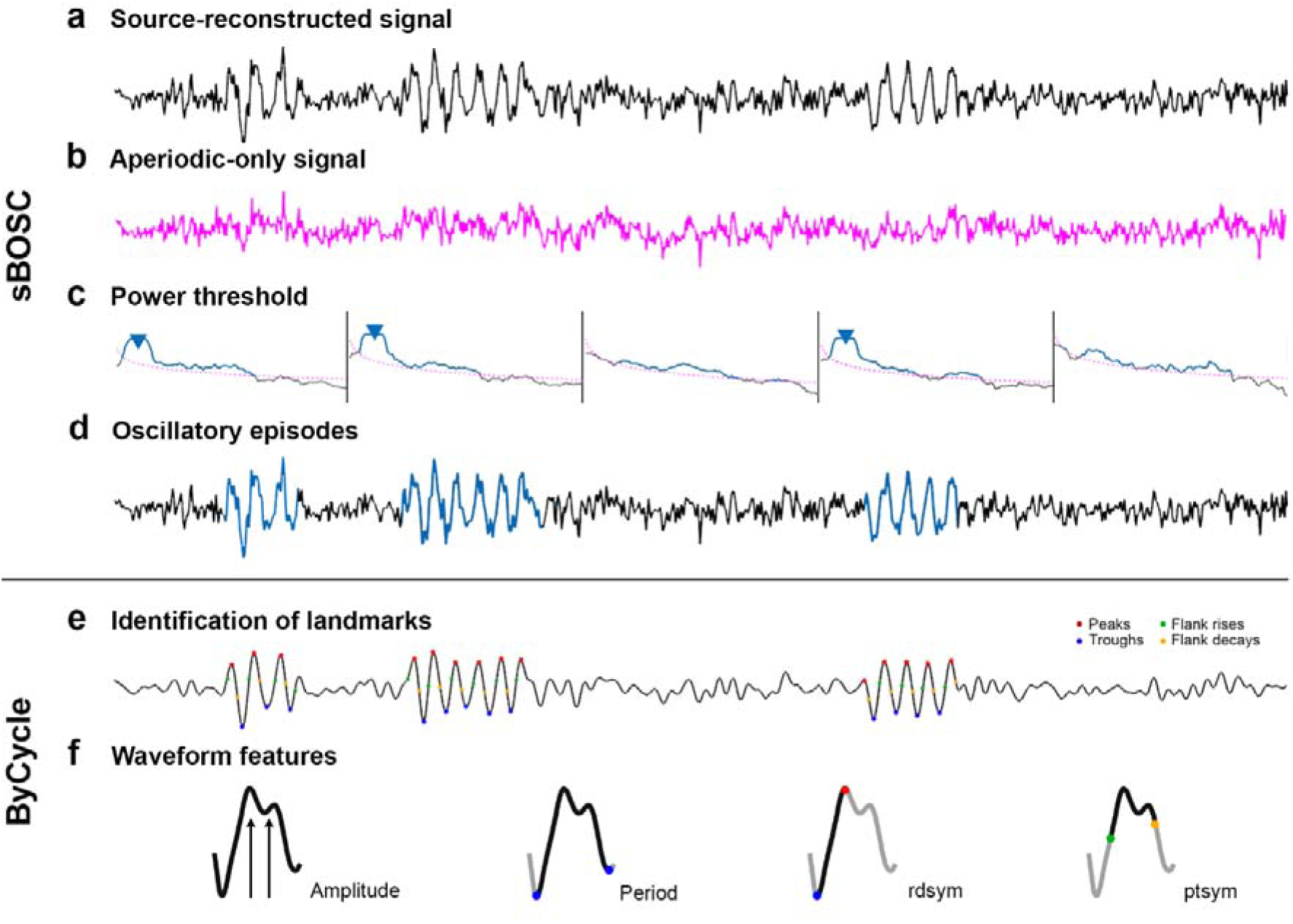
Overview of the two-step (sBOSC and ByCycle) pipeline applied to resting-state MEG data to extract oscillatory episodes and single-cycle metrics. **(a)** Source-level time series were reconstructed via beamforming. **(b)** The aperiodic component was parameterized (specparam) and transformed back into the time domain to obtain an aperiodic-only signal. **(c)** Power spectra of both the original (black) and aperiodic-only signals were computed. A frequency-specific power threshold (pink) was defined as the 95^th^ percentile of the aperiodic power distribution across voxels. Local maxima in both spectral and spatial domains exceeding this threshold were identified as candidate oscillations (blue triangles). **(d)** Segments satisfying all criteria and persisting for at least three cycles were tagged as oscillatory episodes (blue). **(e)** ByCycle was applied to the previously identified oscillatory episodes to segment individual cycles and extract key morphological landmarks, such as peaks, troughs, and flank midpoints. **(f)** For each cycle, we then computed the instantaneous amplitude, period, and two asymmetry metrics: rise-decay asymmetry (rdsym) and peak-trough asymmetry (ptsym).

Second, the aperiodic component was parameterized using the specparam algorithm (Donoghue et al., 2020) in 20-second time windows. The estimated aperiodic component from each window was then back-projected into the time domain to obtain an aperiodic-only time series (Figure 1b). The root mean square (RMS) of this signal was then used as a correction factor to mitigate the center of the head bias in the previously reconstructed source-level time series.

Third, both the original and the aperiodic-only signals were downsampled to 128 Hz and subjected to spectral decomposition using the short-time Fourier transform (STFT) with a 5-cycle adaptive Hanning-tapered sliding window. Power was computed across 32 logarithmically spaced frequency bins spanning 1.8 to 40.5 Hz. A power threshold was then derived from the aperiodic-only power spectra, defined as the 95^th^ percentile of the aperiodic power distribution across voxels for each frequency. Local maxima in the power spectra exceeding this aperiodic threshold were identified as candidates for oscillatory activity (Donoghue et al., 2022; Gross, 2014; Lopes da Silva, 2013). In addition, we detected spatial local maxima of spectral power within the three-dimensional brain volume to minimize the effects of source leakage (O’Neill et al., 2015) (Figure 1c).

Lastly, we identified oscillatory episodes that satisfied all prior criteria (i.e., exceeding the aperiodic power threshold and exhibiting local maxima in both spectral and spatial domains) and sustained a minimum duration of three cycles. To prevent false negatives from brief power drops, sequential oscillatory bursts with the same frequency occurring within the same voxel region (≤ 1.5 cm radius) were merged if separated by less than half a cycle (Figure 1d).

This procedure yielded a four-dimensional binary matrix of participants × voxels × frequencies × time points, where one denotes the presence of an oscillatory episode at a given time point and zero indicates its absence.

### 2.3. ByCycle

ByCycle is an algorithm designed to detect and parameterize individual cycles of brain oscillations in the time domain, enabling the quantification of single-cycle metrics (Cole & Voytek, 2019). We applied ByCycle to the previously identified matrix of oscillatory episodes obtained with sBOSC, thereby restricting the cycle-based analysis to the brain sources and time segments in which oscillatory episodes were present.

Briefly, the ByCycle algorithm segments the signal into successive cycles and identifies relevant key morphological landmarks, including rise and decay zero-crossings, peaks and troughs, and flank midpoints (Figure 1e). Landmark detection was performed across five canonical frequency bands: delta (2 to 4 Hz), theta (4 to 8 Hz), alpha (8 to 12 Hz), low-beta (12 to 20 Hz), and high-beta (20 to 33 Hz). For each band, a low-pass filter set at four times the mean frequency band of interest was applied prior to landmark identification.

Single-cycle metrics were then extracted to quantify distinct oscillatory waveform features (Figure 1f). The cycle band amplitude is defined as the average instantaneous amplitude of the band-pass filtered signal over the duration of the cycle. The period is the time between two consecutive troughs. Rise-decay asymmetry (rdsym) is defined as the proportion of the cycle spent in the rising phase, while peak-trough asymmetry (ptsym) corresponds to the proportion of the cycle occupied by the peak. In the standard ByCycle algorithm, asymmetry metrics (rdsym and ptsym) are centered at 0.5, where deviations from this midpoint indicate the direction of asymmetry (e.g., a shorter rise than decay). However, to facilitate the identification of highly asymmetric regions across the brain, we rescaled both metrics to a 0–1 range, where 0 represents perfect symmetry and 1 represents maximal asymmetry, irrespective of direction.

### 2.4. Analysis of oscillatory episodes and single-cycle metrics

To characterize the proportion of time each brain region exhibited oscillatory activity throughout the recording, we analyzed the oscillatory episodes previously identified with sBOSC. For each participant, we first quantified the percentage of time each brain voxel contained an oscillatory episode, at any frequency. Thus, averaging across participants and voxels yielded a global estimate of oscillatory prevalence in the brain. We further extended this analysis to individual frequency bands by computing the proportion of oscillatory time within each band, as well as the proportion of time each frequency band co-occurred with other bands.

To assess regional differences in oscillatory prevalence across frequency bands, brain voxels were aggregated into five broad areas (frontal, parietal, temporal, occipital, and medial), each comprising eight regions of interest (ROIs) defined according to the Automated Anatomical Labeling (AAL) atlas. Additionally, oscillatory prevalence was z-scored across voxels and then averaged across all constituent voxels within each area, thereby highlighting characteristic frequencies of each brain region. To statistically compare oscillatory prevalence between frequency bands, we conducted separate repeated-measures ANOVAs for each of the five brain areas, with frequency band as the within-subjects factor. Significant effects were followed by post-hoc pairwise comparisons between frequency bands, applying Bonferroni correction to control for multiple comparisons.

In addition to oscillatory prevalence, we extracted metrics to characterize the oscillatory episodes, such as duration and power. As a first approximation, we computed the mean duration of oscillatory episodes across all voxels for each frequency band. To evaluate whether the average number of cycles differed across frequency bands, we conducted a repeated-measures ANOVA with frequency band as the within-subjects factor. Significant main effects were further examined using Bonferroni-corrected post-hoc pairwise comparisons.

Subsequently, to identify specific brain regions exhibiting longer- or shorter-lasting oscillatory activity, we spatially normalized episode duration and applied one-sample t-tests against zero (i.e., the global mean) for every voxel. A non-parametric permutation test was implemented to correct for multiple comparisons, by randomly inverting the signs across participants and retaining the maximum absolute t-statistic across all voxels for a total of 1000 permutations. Significance was established by thresholding the observed t-values at the 95^th^ percentile of the empirical distribution. To disambiguate duration from power, we applied a similar procedure to oscillatory episode power and computed voxel-wise Pearson correlation coefficients between the spatially normalized duration and power values. The statistical significance was evaluated using the aforementioned permutation-based approach.

Lastly, to analyze individual cycle features (amplitude, period, rdsym, and ptsym), we conducted a cycle-by-cycle analysis for each canonical frequency band. The average value for each cycle feature was computed, resulting in a single representative cycle estimate within each participant, brain voxel, and frequency band. Subsequently, cycle features were z-score normalized across voxels within each participant, resulting in a spatial mean of zero. We then performed voxel-wise one-sample t-tests against zero to assess whether each feature significantly deviated from zero, with correction for multiple comparisons using the same permutation-based pipeline described above.

## 3. Results

Oscillatory episodes detected with sBOSC were characterized in terms of their prevalence, duration, and power. We first quantified the proportion of time the brain spent in an oscillatory state, and then extended this analysis to canonical frequency bands and brain regions. Subsequently, the ByCycle algorithm was applied to identify individual cycles within the previously detected oscillatory episodes. This approach enabled the estimation of regional and frequency-band differences in single-cycle metrics, including instantaneous amplitude, period, and waveform asymmetry.

### 3.1. Oscillatory prevalence

To quantify the average temporal prevalence of rhythmic activity in the brain, we first calculated the proportion of total recording time during which oscillatory episodes were present for each participant and voxel, collapsing across all frequency bands. This resulted in a global estimate of 25.14% (± 6.50% across participants, ± 12.83% across voxels; mean ± SD) (Figure 2).

**Figure 2.**
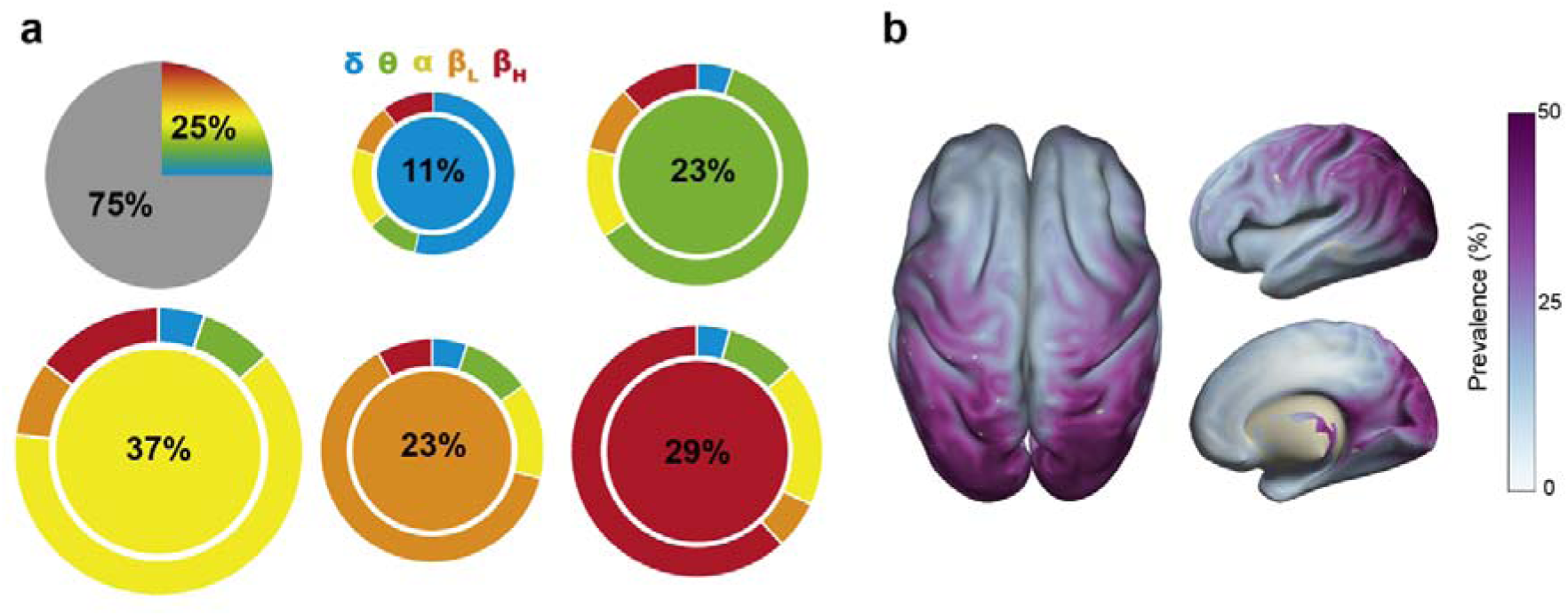
Whole-brain oscillatory prevalence. **(a)** The top-left pie chart shows the overall percentage of oscillatory time in the brain, independent of frequency band. Bubble plots represent the proportion of total oscillatory time during which the brain oscillates within each canonical frequency band (delta in blue, theta in green, alpha in yellow, low-beta in orange, and high-beta in red). Bubble size reflects this proportion. Outer rings indicate the percentage of time each frequency band oscillates either exclusively at its own frequency or in co-occurrence with other bands. **(b)** Superior, lateral, and medial views of the brain illustrate voxel-wise oscillatory prevalence (percentage of time) irrespective of frequency band.

When examined by frequency band, delta oscillations accounted for 11.27% of all detected oscillatory episodes. Of these, 59.88% occurred exclusively within the delta band, whereas 40.12% were shared with other frequency bands. Theta oscillations comprised 22.83% of total oscillatory activity, with 65.39% occurring exclusively within the theta band and 34.61% shared with other frequency bands. Alpha activity constituted 36.67% of total oscillatory activity, with 67.15% oscillating specifically within the alpha band and 32.85% overlapping with other bands. The low-beta band represented 22.61% of overall oscillatory activity, with 67.05% occurring exclusively within low-beta and 32.95% shared with other frequency bands. Finally, the high-beta band was detected in 28.87% of oscillatory activity, with 65.78% oscillating exclusively within that band and 34.22% shared across the remaining bands. The relative co-occurrence of oscillatory episodes across each frequency band and all other bands is represented in the outer rings of Figure 2a. Due to these co-occurrences, the sum of the individual frequency band oscillatory percentages exceeds 100% of the total detected oscillatory time.

Additionally, to identify which brain regions exhibited higher or lower oscillatory prevalence independent of frequency, we computed the proportion of oscillatory time for each voxel, collapsing across frequency bands. As shown in Figure 2b, oscillatory prevalence was higher in parieto-occipital cortices and lower in medial fronto-temporal regions. The voxels exhibiting the highest oscillatory prevalence were symmetrically distributed within the superior lateral occipital cortex, reaching up to 67%, whereas deeper voxels situated in the middle cingulate cortex displayed markedly lower oscillatory activity, with prevalence values of approximately 3%.

<u>Supplementary Table 1</u> summarizes the group-level regional distribution of oscillatory time, reporting the mean percentage of recording time spent in an oscillatory state for each frequency band and AAL-based ROI. For each ROI, a single representative estimate of oscillatory time was computed per participant and frequency band by taking the median across all voxels within that region. Regions located in the parietal and occipital lobes – particularly the superior and middle occipital gyri, the superior and inferior parietal gyri, the angular gyrus, and the cuneus– displayed the highest oscillatory time, accounting for around 35–50% of the total recording time and up to 20% when considering only alpha oscillations.

To further characterize region-specific activity and mitigate the global predominance of alpha oscillations, data were z-score normalized across voxels (Capilla et al., 2022). Figure3 illustrates the relative deviations in oscillatory prevalence across frequency bands, organized into five broad anatomical divisions (frontal, parietal, temporal, occipital, and medial; see Supplementary Figure S1 for the individual ROI distributions). Repeated-measures ANOVAs revealed significant differences in oscillatory prevalence across frequency bands within all five anatomical areas (Frontal: F_(4, 508)_ = 216.38, *p* < .0001; Parietal: F_(4, 508)_ = 89.41, *p* < .0001; Temporal: F_(4, 508)_ = 183.32, *p* < .0001; Occipital: F_(4, 508)_ = 85.00, *p* < .0001; Medial: F_(4, 508)_ = 184.74, *p* < .0001). Bonferroni-corrected post-hoc comparisons indicated that the frontal area was predominantly characterized by high-beta oscillations, parietal regions were associated with elevated alpha and low-beta activity, temporal regions showed relatively higher delta activity, the occipital cortex was mostly dominated by alpha and low-beta activity, and medial brain regions exhibited an increased proportion of theta oscillations (all *p* < .05, corrected for multiple comparisons).

**Figure 3.**
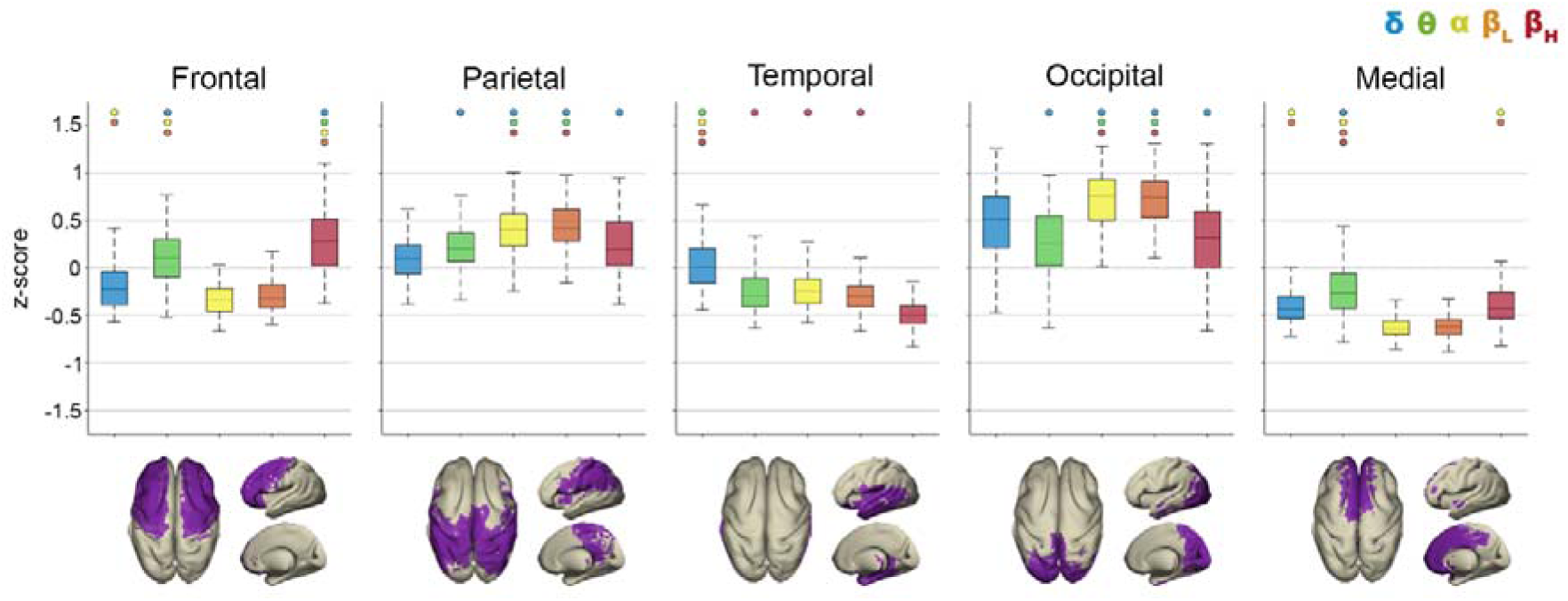
Relative oscillatory prevalence by frequency band and brain area. The figure shows the percentage of oscillatory time z-scored across voxels (grouped into five larger areas: frontal, parietal, temporal, occipital, and medial; bottom row) and frequency band (delta in blue, theta in green, alpha in yellow, low-beta in orange, and high-beta in red). Colored dots above each boxplot indicate that the corresponding frequency band exhibited significantly greater relative oscillatory prevalence than the bands represented by the dot color (*p* < .05, corrected for multiple comparisons).

### 3.2. Duration and power of oscillatory episodes

As an initial characterization of oscillatory episode features, we computed the mean episode duration across all voxels. This yielded an overall mean value of 4.60 ± 0.24 cycles. Specifically, the mean episode duration was 4.07 ± 0.12 cycles (corresponding to 1.35 ± 0.05 seconds) for delta oscillations, 4.31 ± 0.18 cycles (0.72 ± 0.03 seconds) for theta, 5.05 ± 0.34 cycles (0.47 ± 0.04 seconds) for alpha, 4.87 ± 0.30 cycles (0.28 ± 0.01 seconds) for low-beta, and 4.68 ± 0.25 cycles (0.18 ± 0.01 seconds) for high-beta oscillations (Figure 4).

**Figure 4.**
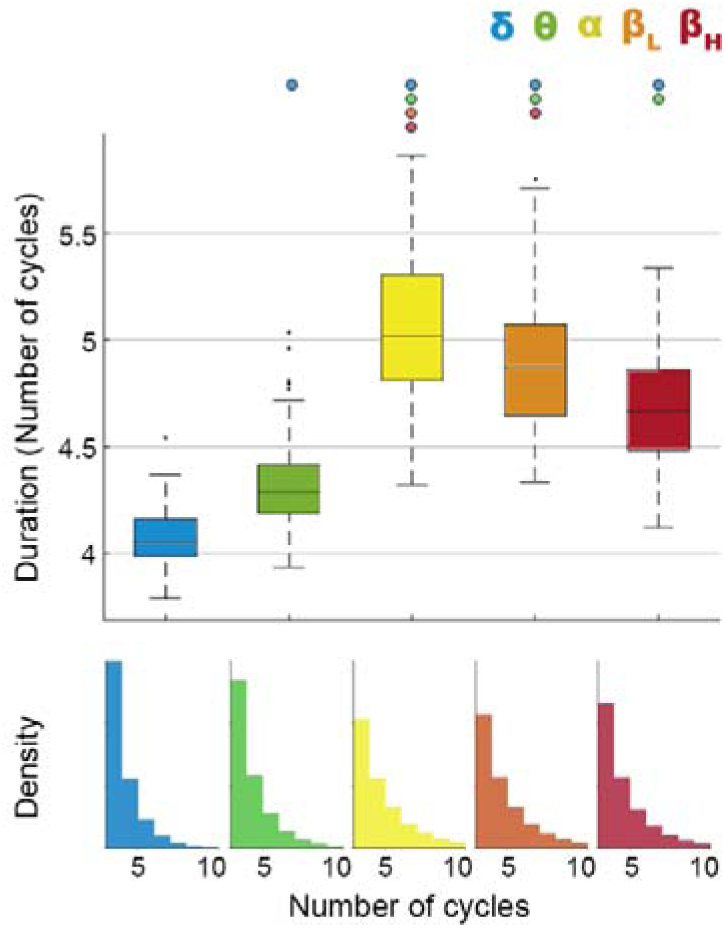
Mean duration of oscillatory episodes by frequency band. Boxplots in the top panel show the global distribution of oscillatory episode duration (in number of cycles) by frequency band. The central line indicates the median, the box represents the interquartile range (IQR = Q3–Q1), and whiskers extend to 1.5×IQR; points beyond this range are shown as outliers. Colored dots above each boxplot indicate that episode duration in the corresponding frequency band was significantly longer than in the bands represented by the dot color (*p* < .05, corrected for multiple comparisons). The bottom panel shows the density distribution of individual oscillatory episode durations across the whole brain, by frequency band.

The repeated-measures ANOVA revealed significant differences across frequency bands (F_(4, 508)_ = 477.04, *p* < .0001), with post-hoc pairwise comparisons confirming differences between all bands (alpha-band episode duration was longer than duration in any other band: t_(127)_ > 6.70, *p* < .0001; low-beta-band duration was longer than duration in high-beta, theta, and delta bands: t_(127)_ > 9.70, *p* < .0001; high-beta-band duration was longer than duration in theta and delta bands: t_(127)_ > 13.81, *p* < .0001; and theta-band duration was longer than delta-band duration: t_(127)_ > 16.86, *p* < .0001; all *p* corrected for multiple comparisons).

Episode duration and power were then z-score normalized across voxels and statistically compared to identify brain regions exhibiting longer-lasting and greater-power oscillatory episodes relative to the whole-brain mean within each frequency band. The spatial distribution of episode duration (Figure 5a) followed a posterior-to-anterior gradient as a function of frequency, with longer-lasting oscillatory episodes predominating in the occipital cortex for the delta and theta bands (∼4 cycles), progressively shifting toward parietal regions at alpha frequencies (∼6 cycles), and toward the frontal cortex in the beta band (∼5 cycles). Notably, the topographic distribution of oscillatory power revealed a distinct frequency-specific organization (Figure 5b). Delta episodes exhibited maximal power in the medial temporal lobe and medial cingulate gyrus, while theta power was highest in the precuneus and superior temporal regions. Alpha power was also maximal in the precuneus, whereas low-beta showed peak power in the sensorimotor cortex. Finally, high-beta oscillations displayed their highest power in frontal regions.

**Figure 5.**
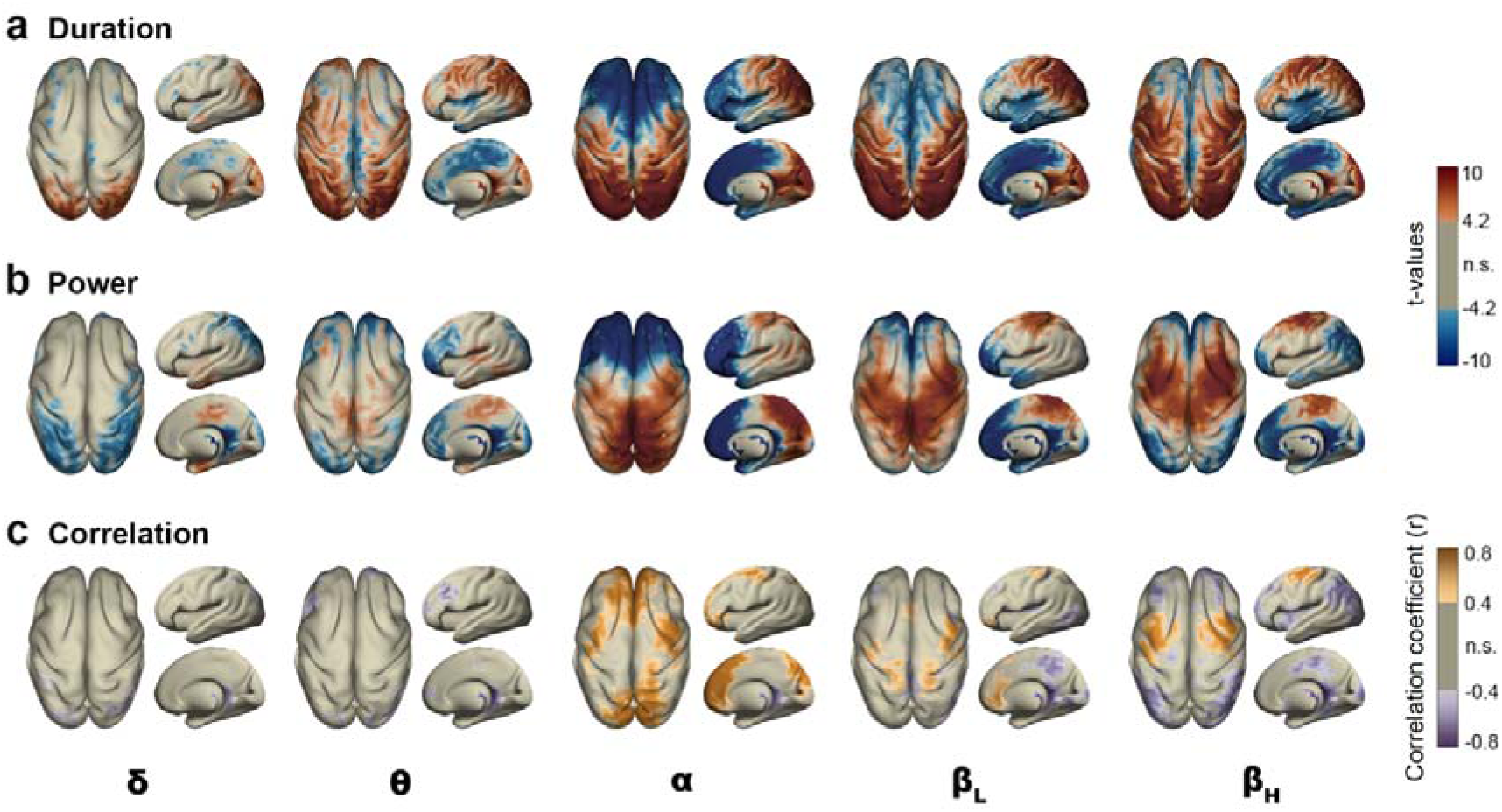
Spatial organization of oscillatory episode duration and power by frequency band. The figure shows superior, lateral, and medial cortical views for each frequency band (columns), displaying **(a)** duration and **(b)** power maps. Red indicates statistically significant positive t-values (above the mean), whereas blue indicates statistically significant negative t-values (below the mean). **(c)** Pearson correlation coefficients between episode duration and power (positive correlations in orange, negative correlations in purple). All maps show only statistically significant results (*p* < .05, corrected for multiple comparisons). Gray areas indicate regions with non-significant (n.s.) effects.

Voxel-wise Pearson correlations between episode duration and power revealed statistically significant positive correlation coefficients in mid-frontal and mid-occipital regions in the alpha band, orbitofrontal cortex in the low-beta band, and lateral-frontal regions within the high-beta band (Figure 5c). For a more detailed analysis of the correlation between duration and power across ROIs, see Supplementary Figure S2.

### 3.3. Cycle-by-cycle estimates of amplitude, period, and asymmetry

Finally, we characterized the waveform morphology of ongoing oscillatory activity across different frequency bands and brain regions by extracting cycle-by-cycle features, such as instantaneous amplitude, period, and asymmetry metrics (rdsym and ptsym). To identify regions with values significantly different from the mean, we z-scored feature values across voxels and conducted one-sample t-tests against zero, with correction for multiple comparisons. Figure 6 shows the brain distribution of the statistical results for each feature and frequency band. Red voxels indicate significantly higher values of a specific feature relative to the global brain average, whereas blue voxels indicate significantly lower values.

**Figure 6.**
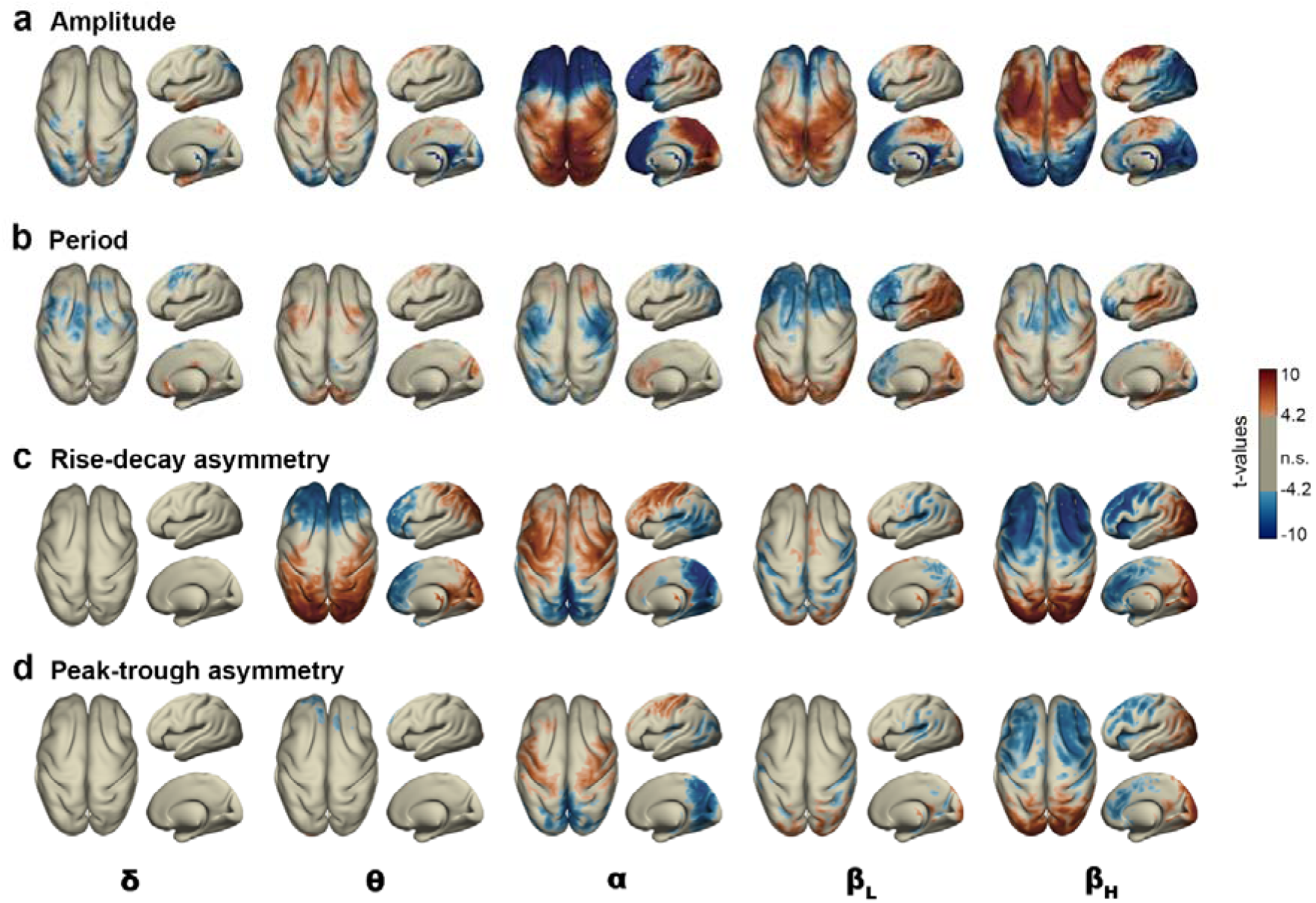
Cycle-by-cycle characterization of amplitude, period, and asymmetry by frequency band. The figure shows superior, lateral, and medial cortical views for each frequency band (columns), displaying t-value maps for **(a)** amplitude, **(b)** period, **(c)** rise-decay asymmetry (rdsym), and **(d)** peak-trough asymmetry (ptsym). Red indicates statistically significant positive t-values (above the mean), whereas blue indicates significant negative t-values (below the mean) (*p* < .05, corrected for multiple comparisons). Gray areas indicate regions with non-significant (n.s.) effects.

The following areas showed significantly higher amplitude across different frequency bands: mid-temporal areas in the delta band, frontal regions in the theta band, occipital and parietal regions in the alpha band, parietal and sensorimotor regions in the low-beta band, and lateral frontal areas in the high-beta band (Figure 6a).

Regarding the period (i.e., the inverse of frequency), faster delta activity (blue) appeared in frontal areas, overlapping with slower theta band frequencies (red). Alpha showed faster oscillations predominantly localized in frontal-lateral areas and slower at mid-frontal regions. Low-beta activity was consistently faster in pre-frontal regions, while slower in occipital and temporal areas. Lastly, high-beta oscillations were faster in frontal and pre-frontal lateral regions while slower in temporal regions (Figure 6b).

Analyses of waveform asymmetry revealed statistically significant different regional and frequency-specific patterns (Figure 6c,d). Delta oscillations showed no significant asymmetry across all evaluated voxels. In the theta band, rise-decay asymmetry was significantly more pronounced in occipital and parietal areas, whereas frontal and medial regions exhibited more symmetric waveforms; however, no significant differences were found in peak-trough asymmetry. Both asymmetry metrics revealed significantly asymmetric oscillations over sensorimotor areas and symmetric at the precuneus and temporal areas within the alpha band. Low-beta waveforms were significantly more asymmetrical in lateral pre-frontal regions and more symmetrical in mid-temporal areas, Finally, high-beta oscillations showed consistent asymmetric patterns in the occipital cortex and symmetric patterns in the frontal cortex across both rise-decay and peak-trough asymmetry indices. Supplementary Figure S3 shows the individual distributions of asymmetry features (rdsym and ptsym) for each ROI and frequency band.

## 4. Discussion

This study aimed to characterize spontaneous whole-brain human oscillatory activity by systematically assessing oscillatory prevalence, episode duration, and waveform morphology across frequency bands and brain regions. By focusing our analysis on oscillatory episodes identified with sBOSC (Stern et al., 2026), we quantified oscillatory time, episode duration, and power across different brain regions and frequencies. We then identified individual cycles within the detected oscillatory episodes using the ByCycle algorithm (Cole & Voytek, 2019). This approach allowed us to further characterize single-cycle waveform properties, such as instantaneous amplitude, period, and asymmetry, rather than relying on the smoothed estimates commonly produced by traditional Fourier-based methods.

### 4.1. Percentage of time spent in an oscillatory state

As a first approximation to characterize the brain’s oscillatory activity, we computed the percentage of time the brain was oscillating, averaged across all voxels and frequency bands. Our results revealed that, on average, the brain spent only ∼25% of the time in an oscillatory state. This finding strongly aligns with recent literature conceptualizing oscillations as transient, intermittent bursts rather than sustained rhythms (Jones, 2016; Tal et al., 2020; Van Ede et al., 2018). However, this does not imply that the brain is silent during the remaining 75% of the time, but rather that individual regions spend the majority of their time engaged in aperiodic activity (Donoghue et al., 2020; He, 2014; He et al., 2010). Notably, as previously reported (Frauscher et al., 2018; Hari et al., 1997), oscillatory activity appears to be confined to a relatively small number of cortical regions, such as the superior and middle occipital gyri or the angular gyrus, which exhibited higher rates of oscillatory time, accounting for approximately 45% of their time.

The observation that rhythmic activity is present for only a fraction of the recording time underscores that power spectral analyses computed over continuous time windows inevitably conflate genuine oscillations with aperiodic activity (Brady & Bardouille, 2022; Cole & Voytek, 2019; Donoghue et al., 2020; Jones, 2016; Kosciessa et al., 2020; Seymour et al., 2022; Stern et al., 2026; Whitten et al., 2011). This conflation is particularly concerning in the analysis of resting-state data, where the absence of a task structure limits experimental control. Furthermore, these considerations extend critically to connectivity analyses, where methods such as phase-amplitude coupling implicitly rely on the assumption of sustained oscillatory activity (Aru et al., 2015; Canolty & Knight, 2010; Colclough et al., 2016).

Each frequency band exhibited a distinct fraction of total oscillatory time, with most oscillations occurring at the alpha (∼37%) and high-beta bands (∼29%). Theta and low-beta frequency bands were present for approximately 23% of the total oscillatory time, while oscillations within the delta frequency band exhibited the lowest prevalence (∼11%), possibly reflecting both the expected attenuation of slow rhythms during alert wakefulness and the inherently reduced sensitivity of MEG to deep medial cortical generators (Gunasekaran et al., 2023). This pattern suggests a marked dominance of faster oscillatory activity during the periods in which the brain is in an oscillatory state, a finding further supported by the high co-occurrence of oscillations within the faster frequency bands.

We also observed regional differences in oscillatory prevalence. Consistent with previous reports, slow delta oscillations were more typical of temporal regions, mid-frontal regions were characterized by theta activity, occipital regions were dominated by alpha and low-beta oscillations, whereas sensorimotor and frontal regions exhibited dominant high-beta activity (Afnan et al., 2023; Capilla et al., 2022; Groppe et al., 2013; Hari et al., 1997; Kalamangalam et al., 2020; Mahjoory et al., 2020, 2020; Mellem et al., 2017; Niso et al., 2016).

### 4.2. Duration and power of oscillatory episodes

Analysis of oscillatory episodes revealed a consistently short temporal structure, with a mean duration of approximately 4.6 cycles. Alpha episodes showed the longest duration (∼5 cycles), followed by low- and high-beta bursts, whereas delta activity exhibited the shortest duration, with an average of ∼4 cycles. This is in line with the prevailing view that oscillatory activity during wakeful, resting state is organized into brief, transient bursts rather than sustained, continuous rhythms (Bullock et al., 2003; Jones, 2016; Seedat et al., 2020; Van Ede et al., 2018; Van Es et al., 2025).

It is important to note that the BOSC framework requires a minimum of three consecutive cycles for episode detection to differentiate oscillatory activity from transient events (Kosciessa et al., 2020; Seymour et al., 2022; Stern et al., 2026; Whitten et al., 2011). This duration criterion could, in principle, be adjusted; however, reducing the minimum number of cycles would inevitably increase the rate of false positive detections. Consequently, we retained this criterion, acknowledging that it constrains the present results to episodes lasting at least three consecutive cycles and may introduce an upward bias in duration estimates by systematically excluding shorter oscillatory events.

As for oscillatory prevalence, we also found regional differences in the duration and power of oscillatory episodes. Occipital regions consistently exhibited the longest episodes across all frequency bands; as frequency increased, this pattern progressively extended to encompass parietal and sensorimotor regions. The regional distribution of power showed a similar frequency-dependent gradient, although there were some notable differences between the episode duration and the power maps, particularly for the slowest frequency bands (delta and theta rhythms). For both delta and theta bands, occipital regions showed the longest episode duration. However, episode power was maximal in mid-temporal and mid-frontal cortices in the delta band, and in parietal and superior temporal cortices in the theta band. Correlation analysis complemented these findings by showing that, although episode duration and power were positively correlated in some brain regions –especially in the alpha band–they were weakly or even negatively correlated in others. Such negative correlations suggest that certain regions tend to exhibit either sustained, low-amplitude oscillatory activity or brief, high-amplitude oscillatory bursts. Whereas most studies have focused on power alone, our findings underscore that both episode duration and power represent dissociable dimensions (Kosciessa et al., 2020; Seedat et al., 2020), and therefore may provide complementary insights into brain oscillatory activity.

### 4.3. Individual cycle waveform features

Beyond characterizing oscillatory episodes in terms of prevalence, duration, and power, we quantified individual cycle features across frequency bands and brain regions using the ByCycle algorithm (Cole & Voytek, 2019). Unlike Fourier-based methods, which produce smoothed spectral estimates, cycle-by-cycle analysis captures instantaneous amplitude and period, as well as waveform asymmetry features, that are obscured by conventional spectral analysis.

Spatially normalized instantaneous amplitude maps revealed a consistent topographic distribution of amplitude across frequency bands, broadly in line with the spatial distribution of power of detected episodes. Delta amplitude was maximal in mid-temporal regions, as well as in the precuneus, while theta amplitude was highest in frontal and medial regions. The topographical distribution of alpha and beta frequency bands amplitude showed the expected posterior-to-anterior gradient (Afnan et al., 2023; Capilla et al., 2022; Groppe et al., 2013; Hari et al., 1997; Kalamangalam et al., 2020; Mahjoory et al., 2020; Mellem et al., 2017; Niso et al., 2016).

The analysis of oscillatory period within each frequency band revealed distinct spatial patterns. Notably, delta oscillations were fastest in the frontal regions where theta oscillations were slowest. Similarly, oscillatory frequency increased systematically from posterior to anterior regions across alpha and beta bands. These findings highlight that oscillatory activity does not conform to discrete spectral categories implied by canonical frequency bands, but instead forms a spectral continuum (Cohen, 2021; Donoghue et al., 2022). Furthermore, distinct rhythms can coexist within a single frequency band at separate brain regions, as is the case with alpha oscillations, which tend to occur over sensorimotor regions at a significantly higher frequency (∼11 Hz) than the global alpha mean –a spectral signature characteristic of the classical mu rhythm (Hari et al., 1997; Tiihonen et al., 1989).

Lastly, the analysis of waveform morphology –rise-decay and peak-trough asymmetry– revealed prominent regional variations across the theta to high-beta frequency ranges. Within the alpha band, sensorimotor regions that exhibited the fastest alpha frequency (i.e., the mu rhythm) also displayed a more pronounced asymmetry across both rise-decay and peak-trough indices. This contrasted with the highly symmetric alpha oscillations observed in the precuneus and temporal cortex. These morphological findings align with the well-documented comb-shaped morphology of the mu rhythm, further supporting its physiological dissociation from posterior alpha oscillations (Bender et al., 2025; Cole et al., 2017; Cole & Voytek, 2017; Hari et al., 1997; Schaworonkow, 2023; Schaworonkow & Nikulin, 2019). Although both asymmetry metrics revealed similar patterns in the alpha and beta frequency bands, they capture distinct morphological features (Cole & Voytek, 2019). This is exemplified by theta band oscillations, where rise-decay –but not peak-trough–asymmetry showed significantly greater symmetry over anterior regions and greater asymmetry over posterior regions. The successful capture of the well-established sinusoidal nature of alpha oscillations in the precuneus and the asymmetry of the mu rhythm in sensorimotor regions validates these metrics and opens the door to uncovering previously unknown waveform morphologies across other frequency bands and cortical regions.

## 5. Conclusion

In this study, we provide a comprehensive characterization of intrinsic neural oscillations in humans. Moving beyond conventional Fourier-based estimates of spectral power, we systematically assessed the proportion of time the wakeful brain engages in oscillatory activity, as well as a range of metrics characterizing oscillatory episodes and waveform morphology. We found that, on average, the brain engages in oscillatory activity for only a small fraction of the time (∼25%) and that oscillatory episodes persist for approximately 4.6 cycles, supporting the view that neural oscillations are expressed as transient bursts rather than sustained rhythms. Interestingly, episode duration and power, along with single-cycle waveform features, exhibited distinct regional distributions across frequency bands. These findings suggest that these metrics may capture complementary dimensions of oscillatory dynamics and provide novel insights into the spatial, temporal, and spectral organization of neural activity in the healthy human brain.

## Author Contributions

**Enrique Stern**: Conceptualization, Data curation, Formal analysis, Methodology, Software, Visualization, Writing – original draft preparation, Writing – review & editing; **Almudena Capilla**: Conceptualization, Data curation, Funding acquisition, Methodology, Supervision, Writing – original draft preparation, Writing – review and editing.

## Declaration of Competing Interest

The authors declare no competing interests.

## Acknowledgements

We are thankful to the contributors of Open MEG Archive (OMEGA) for providing the publicly available dataset used in this study. This work was supported by Ministerio de Ciencia, Innovación y Universidades / Agencia Estatal de Investigación, Spain / FEDER/FSE+, UE (MCIN/AEI/10.13039/501100011033/FEDER/FSE+, UE; grants PRE2022-101613 to ES, and PID2021-125841NB-I00 and PID2024-161032NB-I00 to AC).

## Notes

### Competing Interest Statement

The authors have declared no competing interest.

