## Supplemental material for "Beyond power: A large-scale characterization of intrinsic brain oscillatory activity"

**Supplementary Table 1.** Mean ± standard deviation of the percentage of oscillatory time by ROI and frequency band.

| Region | Total | | | δ | | | θ | | | α | | | $\beta_L$ | | | $\beta_H$ | | |
|---|---|---|---|---|---|---|---|---|---|---|---|---|---|---|---|---|---|---|
| **Frontal** | | | | | | | | | | | | | | | | | | |
| *Precentral gyr.* | 27.35 | ± | 12.01 | 1.83 | ± | 1.27 | 5.87 | ± | 3.28 | 7.96 | ± | 5.5 | 4.7 | ± | 3.8 | 9.9 | ± | 6.67 |
| *Sup. frontal gyr. (dorsolat)* | 18.23 | ± | 8.49 | 1.65 | ± | 1.29 | 5.84 | ± | 3.54 | 3.08 | ± | 2.22 | 2.39 | ± | 1.95 | 6.19 | ± | 4 |
| *Sup. frontal gyr. (orb)* | 11.44 | ± | 6.7 | 1.22 | ± | 1.12 | 2.92 | ± | 2.36 | 2.34 | ± | 2.01 | 1.64 | ± | 1.28 | 3.69 | ± | 2.93 |
| *Mid. frontal gyr.* | 14.9 | ± | 8.46 | 1.31 | ± | 1.11 | 4.24 | ± | 2.95 | 2.55 | ± | 2.22 | 1.96 | ± | 1.77 | 5.42 | ± | 4.25 |
| *Mid. frontal gyr. (orb)* | 19.19 | ± | 9.44 | 1.93 | ± | 1.69 | 4.68 | ± | 3.29 | 3.62 | ± | 3.03 | 2.86 | ± | 2.21 | 7.62 | ± | 4.75 |
| *Inf. frontal gyr. (oper)* | 22.37 | ± | 11.46 | 2.15 | ± | 1.71 | 5.69 | ± | 3.64 | 4.87 | ± | 3.64 | 3.47 | ± | 2.83 | 8.28 | ± | 6.35 |
| *Inf. frontal gyr. (tri)* | 20.4 | ± | 10.86 | 1.98 | ± | 1.73 | 5.09 | ± | 3.68 | 4.05 | ± | 3.3 | 2.96 | ± | 2.32 | 8.13 | ± | 5.99 |
| *Inf. frontal gyr. (orb)* | 19.65 | ± | 10.94 | 2.01 | ± | 1.69 | 4.47 | ± | 3.24 | 4.57 | ± | 3.84 | 3 | ± | 2.47 | 7.25 | ± | 5.61 |
| **Parietal** | | | | | | | | | | | | | | | | | | |
| *Rolandic operculum* | 17.13 | ± | 8.08 | 1.69 | ± | 1.2 | 3.59 | ± | 2.05 | 5.47 | ± | 3.68 | 3.14 | ± | 2.1 | 3.65 | ± | 2.58 |
| *Postcentral gyr.* | 26.76 | ± | 11.97 | 1.63 | ± | 1.15 | 4.52 | ± | 2.35 | 9.92 | ± | 6.62 | 5.98 | ± | 4.09 | 8.24 | ± | 6.14 |
| *Sup. parietal gyr.* | 37.74 | ± | 15.95 | 2.79 | ± | 2 | 6.41 | ± | 3.55 | 15.41 | ± | 9.58 | 9.86 | ± | 6.03 | 11.32 | ± | 7.71 |
| *Inf. parietal gyr.* | 36.2 | ± | 14.84 | 3.12 | ± | 2.16 | 7.18 | ± | 3.81 | 13.8 | ± | 8.68 | 9.3 | ± | 5.7 | 10.06 | ± | 6.76 |
| *Supramarginal gyr.* | 28.78 | ± | 12.4 | 2.32 | ± | 1.6 | 5.72 | ± | 2.95 | 10.58 | ± | 6.68 | 7.05 | ± | 4.49 | 6.88 | ± | 5.48 |
| *Angular gyr.* | 42.21 | ± | 16.34 | 4.44 | ± | 3.13 | 8.63 | ± | 4.86 | 17.75 | ± | 10.31 | 11.88 | ± | 6.81 | 9.81 | ± | 6.93 |
| *Posterior cingulate gyr.* | 29.32 | ± | 16.08 | 3.51 | ± | 2.58 | 5.68 | ± | 4.28 | 11.54 | ± | 8.15 | 6.87 | ± | 5.18 | 6.29 | ± | 4.61 |
| *Paracentral lobule* | 14.41 | ± | 9.97 | 1.01 | ± | 0.89 | 2.82 | ± | 2.28 | 4.48 | ± | 3.86 | 2.38 | ± | 2.3 | 4.07 | ± | 3.95 |
| **Temporal** | | | | | | | | | | | | | | | | | | |
| *Hippocampus* | 18.53 | ± | 9.46 | 3.2 | ± | 2.09 | 4.44 | ± | 3.3 | 6.55 | ± | 4.49 | 3.23 | ± | 2.14 | 2.48 | ± | 1.69 |
| *Parahippocampal gyr.* | 13.27 | ± | 6.91 | 2.29 | ± | 1.75 | 3.19 | ± | 2.85 | 4.05 | ± | 2.92 | 1.9 | ± | 1.17 | 2.16 | ± | 1.26 |
| *Heschl gyr.* | 14.42 | ± | 8.62 | 1.57 | ± | 1.35 | 2.89 | ± | 2.04 | 5.46 | ± | 4.51 | 2.69 | ± | 2.01 | 2.29 | ± | 2.16 |
| *Sup. temporal gyr.* | 21.25 | ± | 10.04 | 2.7 | ± | 1.82 | 4.46 | ± | 2.76 | 8.42 | ± | 6.02 | 4.42 | ± | 3.09 | 2.88 | ± | 2.32 |
| *Mid. temporal gyr.* | 19.97 | ± | 9.23 | 2.41 | ± | 1.93 | 4.1 | ± | 2.86 | 7.45 | ± | 5.23 | 4.49 | ± | 2.8 | 3.23 | ± | 2.39 |
| *Inf. temporal gyr.* | 14.2 | ± | 7.54 | 2.03 | ± | 1.51 | 2.98 | ± | 2.26 | 4.79 | ± | 3.7 | 2.73 | ± | 2.02 | 2.08 | ± | 1.56 |
| *Temporal pole (Mid.)* | 11.45 | ± | 6.41 | 1.53 | ± | 1.23 | 2.45 | ± | 1.84 | 3.14 | ± | 2.55 | 1.73 | ± | 1.29 | 2.59 | ± | 1.61 |
| *Temporal pole (sup)* | 19.43 | ± | 10.01 | 2.42 | ± | 1.79 | 4.33 | ± | 2.91 | 5.75 | ± | 4.61 | 3.28 | ± | 2.26 | 5.2 | ± | 3.83 |
| **Occipital** | | | | | | | | | | | | | | | | | | |
| *Calcarine cortex* | 33.72 | ± | 13.56 | 3.46 | ± | 2.05 | 5.8 | ± | 3.75 | 14.07 | ± | 7.91 | 8.33 | ± | 5.07 | 7.08 | ± | 5.65 |
| *Cuneus* | 36.2 | ± | 19.57 | 3.83 | ± | 3.12 | 6.22 | ± | 4.7 | 17.27 | ± | 12.5 | 9.18 | ± | 7.06 | 7.75 | ± | 7.73 |
| *Precuneus* | 19.77 | ± | 14.27 | 1.67 | ± | 1.44 | 2.81 | ± | 2.44 | 9.28 | ± | 8.31 | 4.04 | ± | 3.84 | 4.39 | ± | 4.97 |
| *Lingual gyr.* | 25.01 | ± | 11.04 | 2.43 | ± | 1.67 | 3.82 | ± | 2.52 | 9.78 | ± | 5.52 | 5.97 | ± | 3.98 | 5.52 | ± | 4.37 |
| *Sup. occipital gyr.* | 48.89 | ± | 16.9 | 4.98 | ± | 3.23 | 8.82 | ± | 4.72 | 22.1 | ± | 11.66 | 13.72 | ± | 7.6 | 13.08 | ± | 9.12 |
| *Mid. occipital gyr.* | 44.33 | ± | 17.69 | 4.93 | ± | 3.62 | 8.14 | ± | 5.17 | 18.88 | ± | 11.05 | 12.23 | ± | 7.03 | 11.78 | ± | 8.36 |
| *Inf. occipital gyr.* | 29.21 | ± | 13.62 | 2.66 | ± | 2.37 | 4.49 | ± | 3.33 | 11.58 | ± | 7.97 | 7.19 | ± | 4.44 | 7.11 | ± | 5.24 |
| *Fusiform gyr.* | 9.5 | ± | 5.73 | 1.61 | ± | 1.24 | 2.22 | ± | 2.07 | 2.95 | ± | 2.37 | 1.34 | ± | 1.06 | 1.37 | ± | 0.82 |
| **Medial** | | | | | | | | | | | | | | | | | | |
| *Supplementary motor area* | 14.49 | ± | 9.12 | 1.3 | ± | 1.28 | 5.1 | ± | 3.89 | 2.74 | ± | 2.11 | 1.61 | ± | 1.87 | 4.58 | ± | 4.05 |
| *Olfactory cortex* | 12.86 | ± | 8.68 | 2.15 | ± | 1.76 | 2.97 | ± | 2.37 | 3.97 | ± | 3.81 | 1.76 | ± | 1.99 | 2.63 | ± | 2.84 |
| *Sup. frontal gyr. (med)* | 16 | ± | 8.71 | 1.56 | ± | 1.5 | 5.7 | ± | 3.96 | 2.65 | ± | 2.07 | 2.12 | ± | 1.69 | 4.95 | ± | 3.31 |
| *Sup. frontal gyr. (med orb)* | 8.1 | ± | 6.28 | 0.89 | ± | 0.98 | 2.2 | ± | 1.78 | 1.5 | ± | 1.72 | 1.06 | ± | 1.01 | 2.48 | ± | 2.93 |
| *Rectus gyr.* | 9.03 | ± | 6.72 | 1.26 | ± | 1.2 | 2.24 | ± | 2.08 | 2.03 | ± | 2.37 | 1.2 | ± | 1.21 | 2.62 | ± | 2.7 |
| *Insula* | 20.64 | ± | 9.77 | 2.44 | ± | 1.6 | 5.44 | ± | 3.2 | 4.92 | ± | 3.4 | 3.13 | ± | 2.39 | 5.47 | ± | 4.28 |
| *Anterior cingulate gyr.* | 11.53 | ± | 7.65 | 1.21 | ± | 1.3 | 3.56 | ± | 2.65 | 2.44 | ± | 2.25 | 1.52 | ± | 1.37 | 3.34 | ± | 3.28 |
| *Mid. cingulate gyr.* | 6.19 | ± | 4.88 | 0.8 | ± | 0.81 | 1.58 | ± | 1.62 | 1.4 | ± | 1.19 | 0.77 | ± | 0.82 | 1.47 | ± | 1.43 |

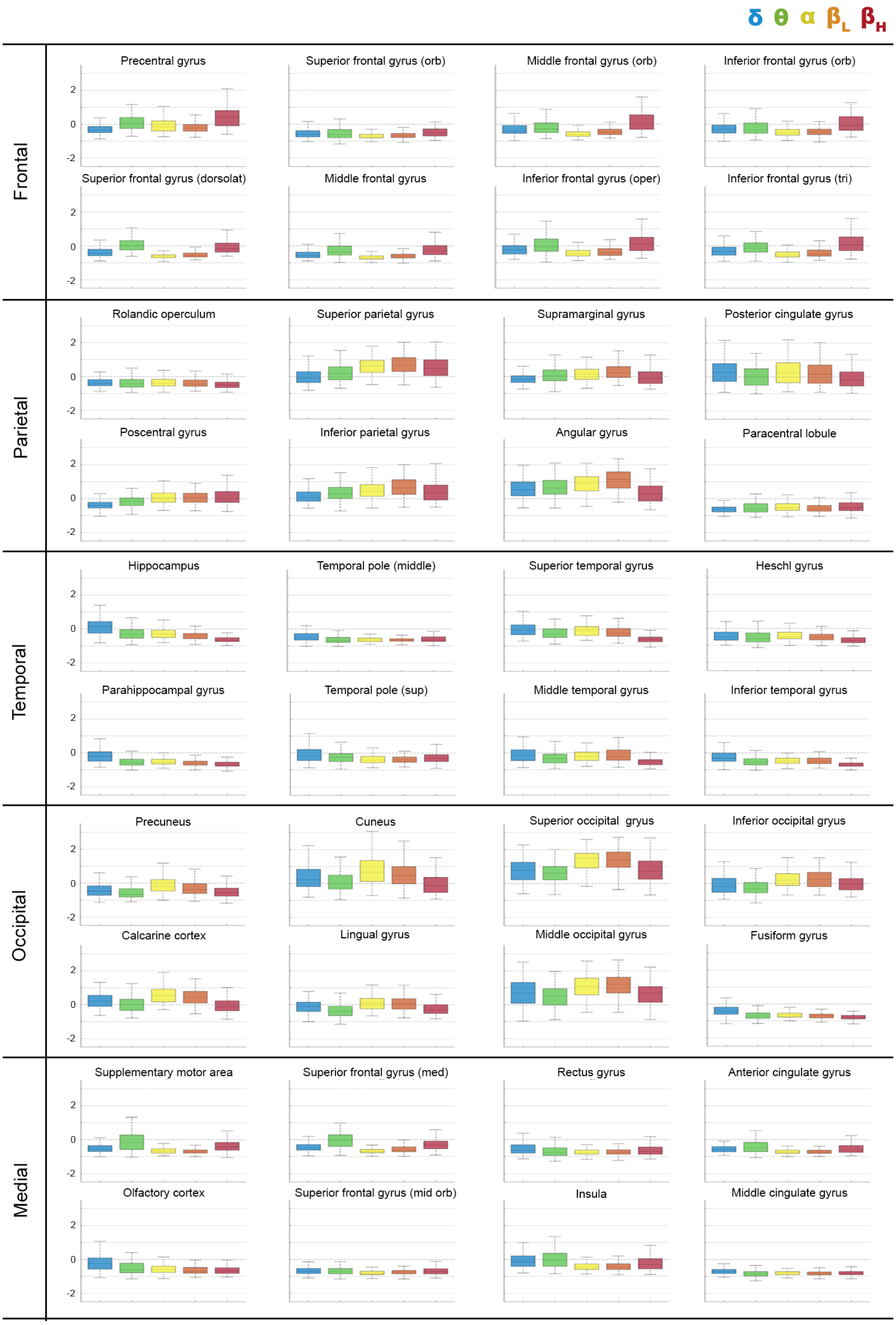


**Supplementary Figure S1. Relative oscillatory prevalence by frequency band and ROI.** The figure shows the distributions of percentage oscillatory time z-scored across voxels for each ROI and frequency band, grouped into five larger areas (frontal, parietal, temporal, occipital, and medial).

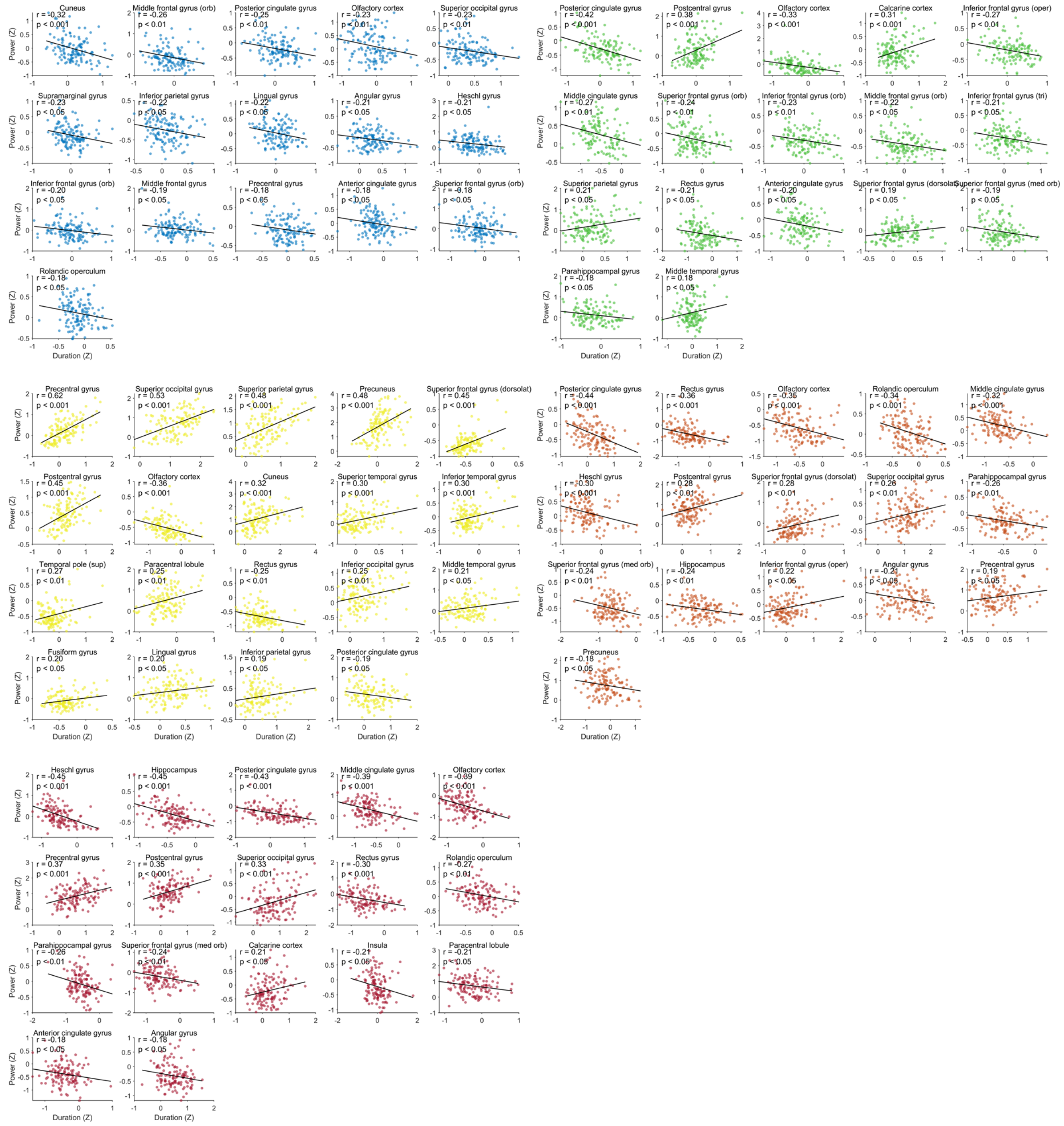


**Supplementary Figure S2. Correlation between episode power and duration (in number of cycles) across ROIs and frequency bands.** ROIs with statistically significant correlations between power and duration are shown ($p < .05$, corrected for multiple comparisons).

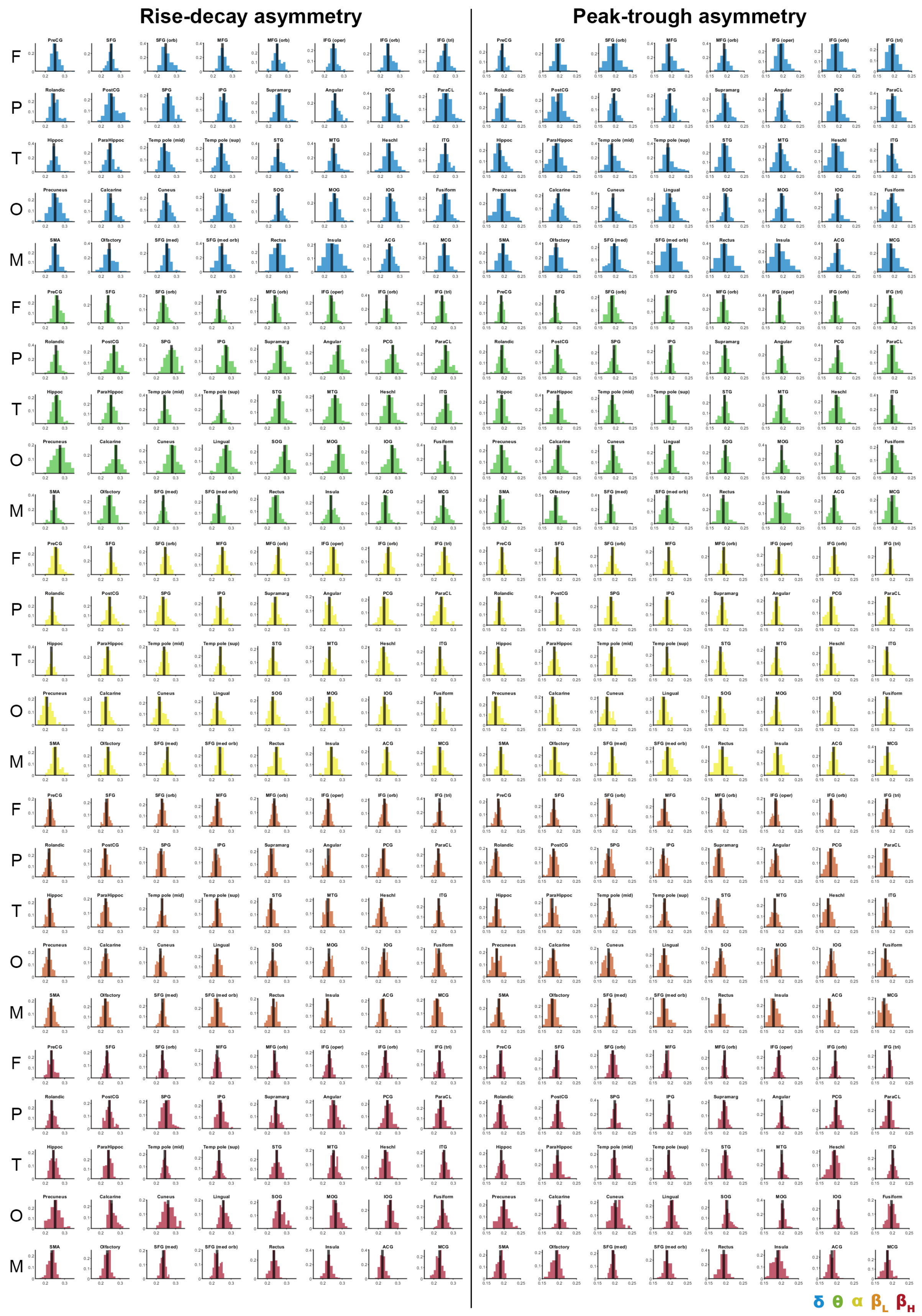


**Supplementary Figure S3. Individual distributions of asymmetry features –rise-decay and peak-trough asymmetry– across ROIs and frequency bands.** Rise-decay asymmetry (left) and peak-trough asymmetry (right) distributions, grouped into five larger areas (F: frontal, P: parietal, T: temporal, O: occipital, M: medial). The black line in each distribution represents the mean value. Note that some ROIs display markedly distinct asymmetry distributions.
